# Structure and functional analyses of Vaccinia virus entry-fusion complex component J5 protein

**DOI:** 10.1101/2025.11.13.688282

**Authors:** Hsiao-Jung Chiu, Louise Tzung-Harn Hsieh, Kathleen Joyce Carillo, Der-Lii Tzou, Wen Chang

**Affiliations:** Molecular and Cell Biology, Taiwan International Graduate Program, Academia Sinica and Graduate Institute of Life Sciences, National Defense Medical Center, Taipei, Taiwan; Institute of Molecular Biology, Academia Sinica, Taipei, Taiwan; Institute of Chemistry, Academia Sinica, Taipei, Taiwan

**Author notes:** These authors contributed equally to this article (Hsiao-Jung Chiu, Louise Tzung-Harn Hsieh, Kathleen Joyce Carillo). Corresponding author: correspondence should be addressed to: Wen Chang; Der-Lii Tzou.

## Abstract

Vaccinia virus enters host cells through an 11-protein entry fusion complex (EFC) that operates by a mechanism distinct from those of canonical viral fusion systems. Understanding how this multiprotein complex mediates membrane fusion is crucial for elucidating poxvirus entry and identifying potential antiviral targets. Here, we determined the NMR structure of a truncated J5 protein, (J5 ^2-68^). To further dissect the functional determinants of J5, we generated recombinant vaccinia viruses expressing various J5 mutants, including substitutions in conserved residues, alterations of exposed charged residues, and chimeric constructs between vaccinia J5 and its orthologous AMV232 gene from an entomopoxvirus. Functional analyses revealed that residues 90-110 and the conserved P^38^YYCWY^43^ motif are indispensable for maintaining EFC integrity and promoting membrane fusion. Together, we define the structural and functional elements of J5 that are essential for poxvirus entry and advance our understanding of the unique membrane fusion mechanism employed by poxviruses.

**Importance:** Vaccinia virus enters host cells through an eleven-protein entry fusion complex (EFC) that is mechanistically distinct from canonical viral fusion systems. Understanding how this multiprotein machinery mediates membrane fusion is essential for elucidating poxvirus entry mechanisms and for developing antiviral strategies. Here, we determined the NMR structure of the J5 ectodomain and generated a series of recombinant vaccinia viruses carrying J5 mutations. Functional analyses identified two regions, residues 90-110 and the conserved P^38^YYCWY^43^ motif, as essential for EFC stability and membrane fusion. These findings provide the first structure-function framework for J5, a core component of the EFC, and reveal key determinants required for complex integrity and viral entry.

## Introduction

Vaccinia virus (VacV) is a prototypic orthopoxvirus that composed of a linear, double-stranded DNA genome with 194.7 kilobase pairs encoding more than 200 open reading frames (1, 2). As the high sequence conservation and similar biological characteristics, VacV has long served as a model system for studying poxvirus molecular biology and virus-host interactions. Historically, vaccinia virus was used as the live vaccine that led to the global eradication of smallpox caused by variola virus. More recently, it has also been employed as a vaccine platform to prevent infection by emerging orthopoxviruses such as monkeypox virus (mpox) during recent outbreaks. (3, 4).

VacV has a broad host range and cytoplasmic life cycle (3); upon infection and subsequent core uncoating, viral factories form at perinuclear regions and further assemble into endoplasmic reticulum-derived single membrane wrapped mature virus (MV) (5, 6). Such intracellular, predominant infectious particles may further pack with Golgi membrane and egress as double-membrane extracellular virus (EV) (7). Although MVs and EVs mediate distinct types of transmission and enter host cells via different approaches (8–10), at post-binding stage, membrane fusion between host endosome and viral particles require an identical multiprotein (11), highly-conserved entry-fusion complex (EFC). The EFC contains 11 proteins: A16 (12), A21 (13), A28(14), F9 (15), G3 (16), G9 (17), H2 (18), J5 (19), L1 (20), L5 (21) and O3 (22). Unlike the vast majority of virus utilizes single viral protein to engage with cellular receptor and further mediate the membrane fusion subsequent to an event resulted into conformational change (23); for poxvirus, at least 4 viral proteins mediate the attachment via glycosaminoglycans or receptors on the cell surface (24–27), in addition, each EFC component is pivotal for the viral infectivity, the formation of the EFC, and membrane fusion activity despite that they may act at different stage of membrane fusion process (28). Recently, several subcomplexes have been identified with protein structure available, which are A16 and G9 (29–31), A28 and H2 (with a dissociation constant at μM range) (32–34) as well as G3 and L5 (35, 36). Another study suggests the EFC protein O3 is in proximal to every other 10 EFC components (37), presumably due to its nature, i.e. a 35-amino acids long α-helix (38). The crystal structures of several EFC components have been reported, except J5 and O3 (31–33, 35, 39–41), yet none of these assessable structures or protein sequences demonstrate homology to the well-characterized fusion peptides or loops that mediated membrane insertion (42) buried in the fusion proteins in class I, II and III (43, 44). Therefore, further investigation of the poxvirus EFC is needed to better understand how these 11-component complex mediate the fusion step during viral entry.

Among these VacV EFC components, there are two structural conserved families, i.e. A16/G9/J5 and F9/L1 (15, 19), classified according to the location of Cysteine residues and the patterns of the disulfide bonding, distribute widely across Poxviridae, non-Poxviridae giant virus and even in prokaryote such as Amoeba and certain mycobacterium (45), suggesting a conserved fusion mechanism during evolution. Since vaccinia L1/F9 serves as a peripheral component in the entry fusion complex (30, 34, 36), we focused on the fusogenic property of the A16/G9/J5 family and how they might dock with the others for EFC assembly. Given that J5, despite having the shortest protein sequence in evolutionary conserved A16/G9/J5 family (19), contains the most structurally conserved elements among the family, and that J5 deletion mutant is impossible to be isolated (46), leading us further characterizing the structural and functional features to address the importance of J5. In this study, we successfully determined an NMR in-solution structure of a truncated form of J5. We also performed mutagenesis analysis and then generated the mutant J5 recombinant virus and identified critical residues of J5 that are essential for triggering membrane fusion and EFC assembly.

## Materials and Methods

### VacV J5 plasmid construction, protein expression and purification

The cDNA encoding the soluble fragment of the vaccinia virus J5 protein (residues 2-68; tJ5) was cloned into the pETDuet-1 vector with an N-terminal His₆-yeast SUMO (Smt3) fusion tag under control of the T7 promoter. The construct was transformed into *E. coli* BL21(DE3) cells for protein expression. For isotopic labeling, the U-^15^N and U-^13^C labeled protein was expressed in M9 minimal media, which contained 1 g/L of ^15^NH_4_Cl (Sigma-Aldrich) as the sole nitrogen source and 1 g/L of U-^13^C_6_ glucose (Cambridge Isotope Laboratories) as the sole carbon source. The media was supplemented with 2 mM of MgSO_4_ (0.24 g/L), 0.1 mM of CaCl_2_ (11.1 mg/L), 1 mg/L of thiamin, and 1 mg/L of biotin. The cells were initially grown overnight in 30 mL of LB medium and then transferred into 1 L of M9 medium after centrifuging to remove the excess LB medium. Protein expression was induced by adding 0.5 mM isopropyl-β-d-thiogalactoside (IPTG) when the optical density of the cells reached 0.6. The cells were incubated for 18 hours at 16 °C and subsequently harvested by centrifugation at 3000 × g for 30 minutes at 4 °C. The cell pellets were resuspended in lysis buffer composed of 20 mM Tris, 50 mM NaCl, 1 mM phenylmethylsulfonyl fluoride (PMSF), 1 mM 1,4-dithiothreitol (DTT) and lysozyme at pH 8.0 until homogeneous. Lysis was achieved by sonication in an ice bath. The lysate was cleared of cell debris by centrifugation (1 hour at 30000 × g at 4 °C) and filtered through a 0.45 μm filter.

The supernatant was incubated overnight at 4°C with 1 mL of Ni^2+^-NTA affinity resin (BioRad) packed into an empty PD10 column. The resin was washed three times with 10 column volumes of 20 mM Tris buffer containing 500 mM NaCl and 20 mM imidazole at pH 8.0. The protein was then eluted using 20 mM Tris, 500 mM NaCl, 500 mM imidazole, and 1 mM DTT at pH 8.0. To remove the His_6_-SUMO tag, Ulp1 protease (47) was used during dialysis with 20 mM Tris and 50 mM NaCl at pH 8.0 overnight at 4°C. The untagged tJ5 protein was subsequently separated using Ni-NTA resin (BioRad). The protein sample was concentrated to 0.5 mM using an Amicon Ultra Centrifugal Filter with a 3 kDa molecular weight cut-off (Millipore). Finally, the concentrated protein was exchanged into an NMR buffer containing 10 mM phosphate (pH 6.5) and 137 mM NaCl and 2.7 mM KCl. The protein concentration was determined by absorbance at 280 nm (ε280 = 13,325 M-1 cm-1), and its purity was confirmed by 15% SDS-PAGE.

### Nuclear magnetic resonance (NMR) spectroscopy and structural determination

All NMR spectra were recorded at 298 K and pH 6.5 using a Bruker AVANCE 600 or 800 MHz spectrometer, equipped with a 5 mm triple resonance TXI cryogenic probe that includes a shielded Z-gradient. Samples of 0.5 mM ^15^N, ^13^C-labeled tJ5 in 1X PBS buffer, with a composition of 90% H_2_O and 10% D_2_O at pH 6.5, were loaded into 5 mm Shigemi NMR tubes for the experiments.

Sequential backbone resonance assignments were achieved through independent connectivity analysis of NHCACB, CBCA(CO)NH, HNCO, and HN(CA)CO experiments. NOE restraints were obtained from 3D ^13^C- and ^15^N-edited NOESY-HSQC spectra with a mixing time of 150 ms. The assignments of backbone residues and NOE resonances were conducted using Topspin 4.3.0 (Bruker) and CARA 1.8.4 (48) software. NOE cross-peaks were categorized as very weak, weak, medium, or strong based on their intensities.

Hydrogen bond restraints were identified by observing the slow exchange of HN signals with the solvent D2O. Dihedral angle restraints (ϕ and Ψ) were predicted from chemical shift data using the TALOS+ web server (49). For the final set of protein structure calculations, 100 structures of tJ5 were generated using the XPLOR-NIH 3.4 software (50, 51) with a standard simulated annealing protocol, starting at high-temperature dynamics (1000 K) and cooling to 100 K. Twenty structures with the lowest energy were chosen for water refinement using the AMPS-NMR web portal (52) and a standard restrained molecular dynamics protocol implemented within the AMBER99SB-ILDN force field (53). This refinement utilized a generalized Born model, an ionic strength of 137 mM NaCl, and a 10 Å TIP3P water box. The final ensemble of 10 tJ5 structures, selected based on conformational energies, had no distance or dihedral angle restraints exceeding 0.5 Å or 5°, respectively. The quality of this final ensemble was assessed and validated using the Protein Structure Validation Suite (PSVS) (54). Protein structure figures were generated using the Chimera 1.14 graphics program (55). Statistics for the resulting structure are summarized in Table 1. The 10 conformers of tJ5 have been deposited in the Protein Data Bank under the PDB ID code 8WT5.

**Table 1.** NMR and refinement statistics for vaccinia virus fusion protein tJ5.

|  | tJ5 (2–68) |
| --- | --- |
| **NMR distance and dihedral constraints** |  |
| Distances constraints |  |
| Total NOE | 836 |
| Intraresidue | 294 |
| Interresidue | 543 |
| Sequence ($\vert i-j\vert = 1$) | 294 |
| Medium-range ($\vert i-j\vert \leq 4$) | 296 |
| Long-range ($\vert i-j\vert \geq 5$) | 119 |
| Dihedral angle restraints | 78 |
| $\phi$ | 39 |
| $\varphi$ | 39 |
| **Hydrogen bond restraints <sup>a</sup>** | 127 |
| **Structure statistics** |  |
| Distance violations per structure |  |
| 0.1-0.2 Å | 6.1 |
| 0.2-0.5 Å | 10.0 |
| > 0.5 Å | 8.3 |
| r.m.s. of distance violation per constraint | 0.1 Å |
| Max. distance violation | 15.5 Å |
| Dihedral angle violations per structure |  |
| 1-5° | 6.3 |
| > 5° | 5.6 |
| r.m.s. of dihedral violation per constraint | 1.0° |
| Max. dihedral angle violations | 19.3° |
| **Ramachandran plot summary <sup>b</sup>** |  |
| Most favored regions | 98% |
| Additionally allowed regions | 1% |
| Generally allowed regions | 1% |
| Disallowed regions | 0% |
| **Averaged r.m.s.d. to the mean structure <sup>c</sup>** |  |
| Backbone | 0.41 Å |
| Heavy atoms | 0.45 Å |

<sup>a</sup> Two distance ranges (NH-O = 1.8-2.4 Å, N-O = 2.8-3.4 Å) were used for the hydrogen bond restraints between the amide and carbonyl group atoms.
<sup>b</sup> Ramachandran plot analysis was performed over residues 2-68 with PROCHECK.
<sup>c</sup> Average r.m.s.d. was calculated from an ensemble of 10 structures over secondary structure elements, including α-helices: residues 3-12, 39-42, 44-47, and 55-64.

### Cells, viruses and reagents

BSC40, CV-1, GFP- and RFP-expressing HeLa cells were cultured in Dulbecco’s modified Eagle’s medium (DMEM) supplemented with 10% fetal bovine serum (HyClone) and 100 units/mL Penicillin and 100 µg/mL Streptomycin (Gibco) as previously described (56). The Western Reserve (WR) strain of vaccinia virus and other recombinant viruses constructed in this study were grown on BSC40 cells. Restriction enzymes were from New England Biolabs. Fixative and staining solutions for electron microscopy analysis were from Electron Microscopy Sciences.

### Bioinformatic analyses of poxviral J5 orthologues

Orthologs of vaccinia J5 protein from representative members of the *Poxviridae* family were retrieved from GenBank. The analyzed accession numbers were: YP_232979.1 (VACV), NP_570485.1 (camelpox virus), NP_619893.1 (cowpox virus), NP_671599.1 (ectromelia virus), NP_536516.1 (monkeypox virus), YP_009282790.1 (skunkpox virus), YP_717406.1 (taterapox virus), NP_042126.1 (variola virus), YP_009281844.1 (volepox virus), NP_039099.1 (fowlpox virus), YP_009177120.1 (turkeypox virus), YP_001293261.1 (goatpox virus), YP_004821430.1 (yokapox virus), QGT49348.1 (crocodilepox virus), YP_008658487.1 (squirrelpox virus), NP_051781.1 (myxoma virus), NP_044029.1 (molluscum contagiosum virus), YP_009480606.1 (sea otterpox virus), NP_957964.1 (bovine papular stomatitis virus), YP_009112794.1 (parapoxvirus), YP_003457360.1 (pseudocowpox virus), NP_957832.1 (orf virus), YP_009268788.1 (pteropox virus), AKR04183.1 (sapmonkeypox virus), NP_938325.1 (yabapox virus), YP_009001515.1 (anomala virus), YP_009408025.1 (eptesipox virus), and NP_065014.1 (Amsacta moorei entomopoxvirus, AMV).

Multiple sequence alignments were generated using MAFFT v7.453 with parameters --maxiterate 1000 --globalpair, and visualized with Jalview v1.8.3. Residue conservation was color-coded: yellow (identical), blue (>0.5 conservation), and green (>0.2 conservation). The protein sequences of VacV J5 and AMV232 were further aligned using Clustal Omega (https://reurl.cc/W8LjQ9). In the alignment, an asterisk (*) denotes identical residues, a colon (:) indicates residues with similar biochemical properties, and a dot (·) marks weakly conserved position.

### Protein Structure Prediction with AlphaFold2

Full-length J5 (YP_232979.1) and AMV232 (NP_065014.1) structures were predicted using AlphaFold2 (https://reurl.cc/96anZj). Sequences were queried using MMseqs2 (mmseqs2_uniref_env) in unpaired/paired mode. For each protein, the predicted model with the highest overall confidence (pLDDT >70) was selected for analysis. The mean pLDDT scores for J5 and AMV232 were 80.7 and 76.5, respectively.

### Residue interaction analysis

The tJ5 structure (PDB: 8WT5) was analyzed using the Residue Interaction Network Generator (RING; https://ring.biocomputingup.it/) to identify intramolecular residue contacts.

### Construction of J5 mutant plasmids and recombinant virus

J5 ORF mutants were synthesized (GenScript Inc.) and cloned into the pUC57 plasmid with *BstZ17*I and *Mfe*I sites at its 5’ and 3’ end, and detailed protein sequences were summarized in the table below.

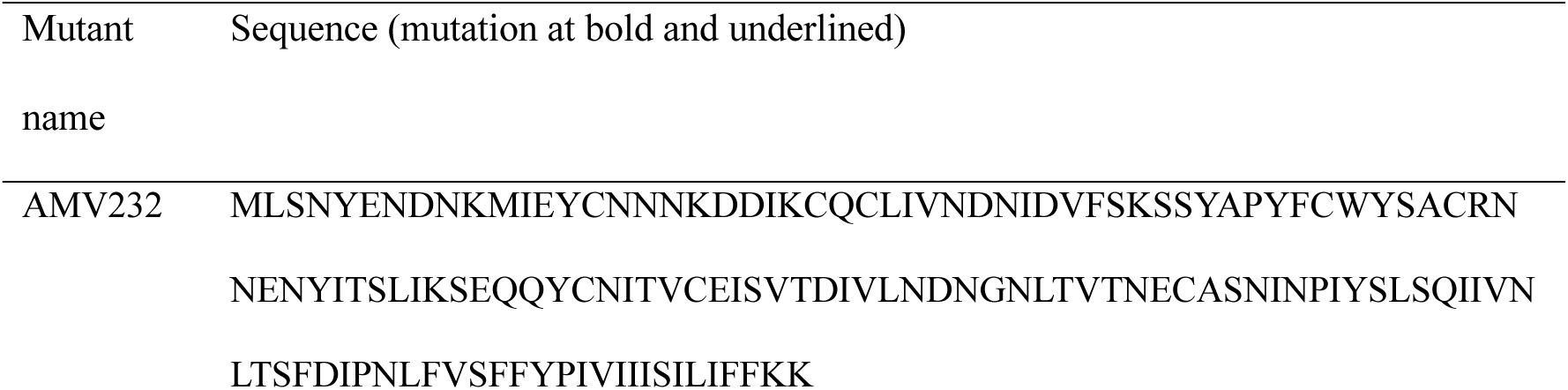
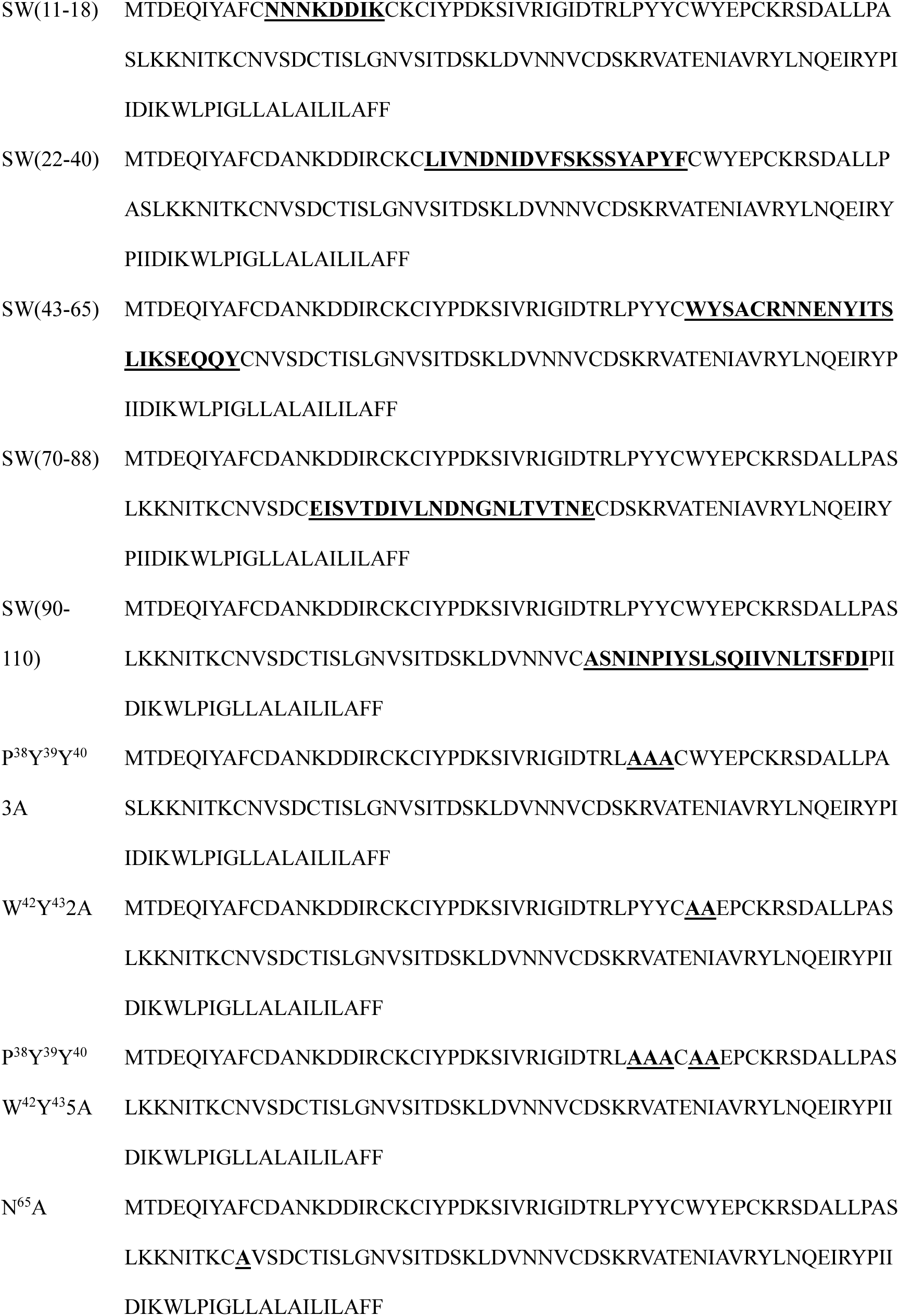
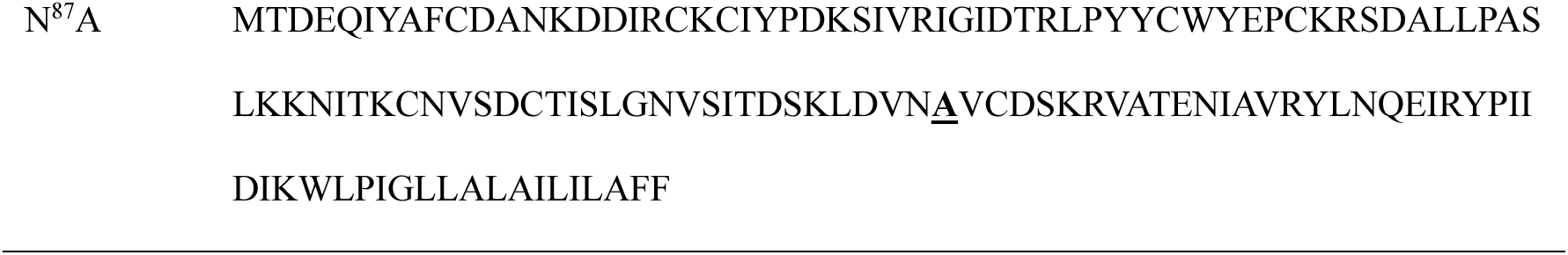

Eco*Gpt* cassette driven by the p7.5k promoter was constructed into a plasmid containing flanking sequences to J5L loci on the viral genome with two mutations as described below. The 1-kb flanking fragment containing partial J6R and J5L promoter sequence was isolated by PCR using WR genomic DNA as template and cloned into pBlueScript KS(-) using primer 5’-TAGGGCGAATTG<u>GGTACC</u>AACGGTGATAGATGTACT-3’ and 5’- GAATTCGATATC<u>AAGCTT</u>GTATACCTTCGGTTCTTT-3’ (the *Kpn*I and *Hind*III site were underlined). Then, a mutation was introduced to destroy a BstZ17I site using primers 5’-AATTCATTGGTTTCTTTGGGAATACTATCTATCCAAAAA CT and 5’-AGTTTTTGGATAGATAGTATTCCCAAAGAAACCAATGAATT-3’. Eco*Gpt* and 1-kb flanking fragment containing J4 ORF and partial J3 were generated by PCR using primers 5’-CAAGCTTGATATCGAATTCAATTGGATCACTAATTCCAAACC-3’ and 5’- TATTTCAGTCGCTGATTAATCTAGCGACCGGAGATTGGCGG-3’, 5’- CCGCCAATCTCCGGTCGCTAGATTAATCAGCGACTGAAATA-3’ and 5’-TGGAGCTCCACCGCGGTGGCGGCCGCCCGTTTCCAGATCAATGGAT-3’ respectively, and subsequently using NEBuilder to assemble with linear pBlueScript-J6. Then, J5 ORF was isolated by PCR using primers 5’-TAT<u>GTATAC</u>AATCAAATTTCCCTTTTTA-3’ and 5’- TAT<u>CAATTG</u>GTAGAGATGAGATAAGAA-3’ or by double digestion of *BstZ17*I and *Mfe*I and subcloned into pBlueScript J6R-gpt-J4R-J3. In several cases, J5 mutants were generated from J5 WT plasmid using QuikChange Lightning site-directed mutagenesis kit (Agilent Technologies) with the primers listed in the table below. The resulting linearized plasmids were separately transfected into WR-GS infected CV-1 cells and then harvested at 24hrs post transfection. Recombinant viruses were subsequently isolated by at least three rounds of plaque purification in 1% agar under *gpt* selection (containing 25 µg/mL mycophenolic acid, 250 µg/mL xanthine and 15µg/mL hypoxanthine) as described in the established protocol (57).

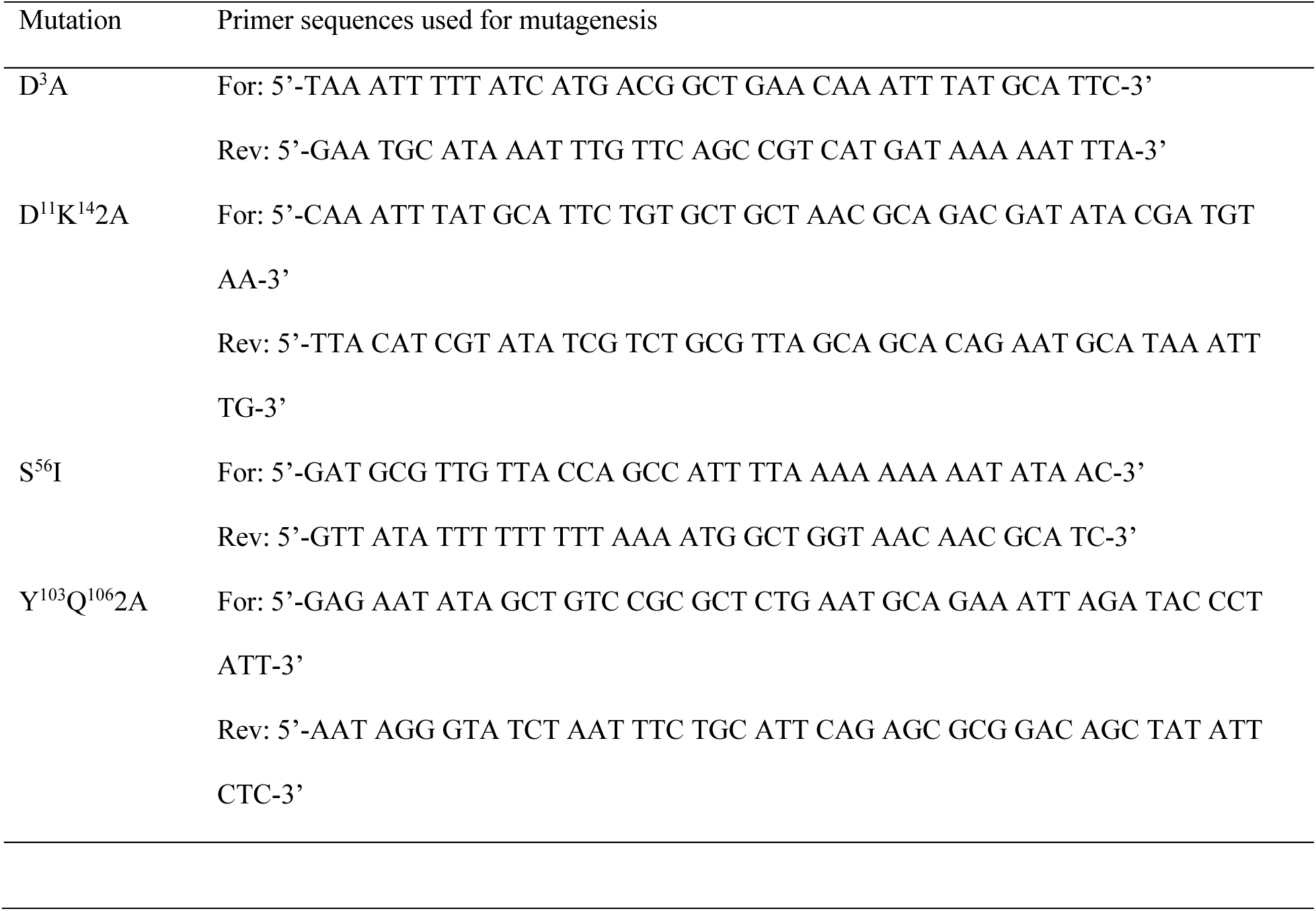

### Determination of J5 mutant virus infectivity

BSC40 cells were seeded in six-well plates and infected with wild-type (WT) or mutant J5 recombinant vaccinia viruses at a multiplicity of infection (MOI) of 5 PFU per cell. After 1 h of adsorption at 37°C, the inoculum was removed and replaced with fresh growth medium. Infected cells were incubated for 24 h post-infection and harvested for plaque assays and immunoblot analysis. All experiments were repeated three times independently.

### MV-triggered cell-cell fusion at acid pH

The MV-mediated cell-cell fusion assays were performed as described previously(32, 33, 56). The HeLa cells expressing EGFP or RFP were mixed at 1:1 ratio and seeded in a 96-well plate. The following day, cells were pretreated with 40 μg/mL cordycepin (Sigma) for an hour and subsequently incubated with J5 recombinant virus infected lysate for an hour at 37 °C. Then, cells were washed, incubated with PBS at either pH 7.4 or 5.0 at 37 °C water bath for 3 minutes, neutralized with growth medium and replaced with fresh medium in the presence of cordycepin. Cell fluorescence was imaged every 30 minutes using ImageXpress Confocal HT.ai High Content Imaging system (Molecular Devices) until 3 hours and cell fusion event was calculated using Fiji software plugin LabKit as a % of the image area of GFP^+^ RFP^+^ cells divided by that of single-fluorescent cells.

### Immunoblot analysis

Protein samples were denatured in SDS sample buffer at 95°C for 10 min and analyzed by immunoblotting. Samples were separated by SDS-polyacrylamide gel electrophoresis (SDS-PAGE) and transferred onto 0.45-µm nitrocellulose membranes (Bio-Rad). The membranes were probed with polyclonal antibodies against A16, A21, A28, D8, F9, G3, G9, H2, J5, L1, and L5 as previously described (33).

### Coimmunoprecipitation

Confluent BSC40 cells were infected with recombinant J5 mutant viruses at a multiplicity of infection (MOI) of 2 PFU per cell and incubated for 24 h at 37°C. Cells were washed with PBS, harvested, and lysed in ice-cold buffer containing 20 mM Tris-HCl (pH 8.0), 200 mM NaCl, 1 mM EDTA, and 0.5% or 1% NP-40 supplemented with protease inhibitors (1 mM phenylmethylsulfonyl fluoride, 2 µg/mL aprotinin, 1 µg/mL leupeptin, and 0.7 µg/mL pepstatin). After centrifugation to remove cell debris, 250 µg of clarified lysate was incubated with protein A-conjugated beads (GE Healthcare) prebound to anti-J5, anti-G9, or anti-A28 antisera. The mixtures were rotated for 16 h at 4°C. Beads were then washed five times with lysis buffer, and bound proteins were analyzed by SDS-PAGE and immunoblotting.

### Electron microscopy analysis

Confluent BSC40 cells were seeded onto poly-L-lysine-coated plastic slides and infected individually with J5 mutant viruses separately at a multiplicity of infection (MOI) of 2 PFU per cell for 24 h. Cells were fixed with glutaraldehyde, postfixed with osmium tetroxide, and contrasted with uranyl acetate. After serial ethanol dehydration, samples were embedded in Epon resin and sectioned for transmission electron microscopy (TEM). Samples were imaged under a Talos L120C transmission electron microscope using a CETA 16 4kx4k CMOS camera (Thermo Scientific).

### Statistical analysis

Statistical analysis was performed using GraphPad Prism v10 (GraphPad Software, San Diego, CA). Data were presented as mean ± standard deviation (SD) Statistical significance was assessed using the two-tailed Student’s *t* test. For comparison with WT controls, the *P* value was adjusted using “p.adjust” function with the false discovery rate (FDR) method in R v4.5.1. Adjusted *P* values of < 0.05 were considered statistically significant. Significance levels are indicated as follows: *, *P*< 0.05; **, *P* < 0.01; ***, *P*< 0.001 and ****, *P* < 0.0001.

## Results

### Nuclear magnetic resonance (NMR) structure of truncated vaccinia virus fusion component J5 (tJ5)

In the NMR structural analysis, we expressed and purified a soluble truncated J5 construct in the pET expression system, denoted as tJ5 (residues 2-68), see Figure 1A-B, which comprises a copy of the J5 ectodomain. The secondary structure of the vaccinia virus fusion protein J5 (residues 1-133) was predicted using AlphaFold2. It revealed six α-helices and inter-connecting loop regions at its N-terminus (residues 1-68), followed by a disordered region and a transmembrane α-helical domain at the C-terminus. By applying ^15^N and ^13^C isotope labeling, we determined the backbone sequential assignments of tJ5 using 3D heteronuclear NMR spectroscopy, as indicated in the ^1^H-^15^N HSQC spectrum (Supplementary Fig. S1A). The 2D HSQC pattern of isotope labelled tJ5 exhibited well-resolved resonances for non-proline residues, suggesting that the protein structure is well-folded. We predicted secondary structure based on the ^13^C chemical shift propensity analysis. The result is in good agreements with structure prediction generated by AlphaFold2 (Supplementary Fig. S1B).

**Figure 1.**
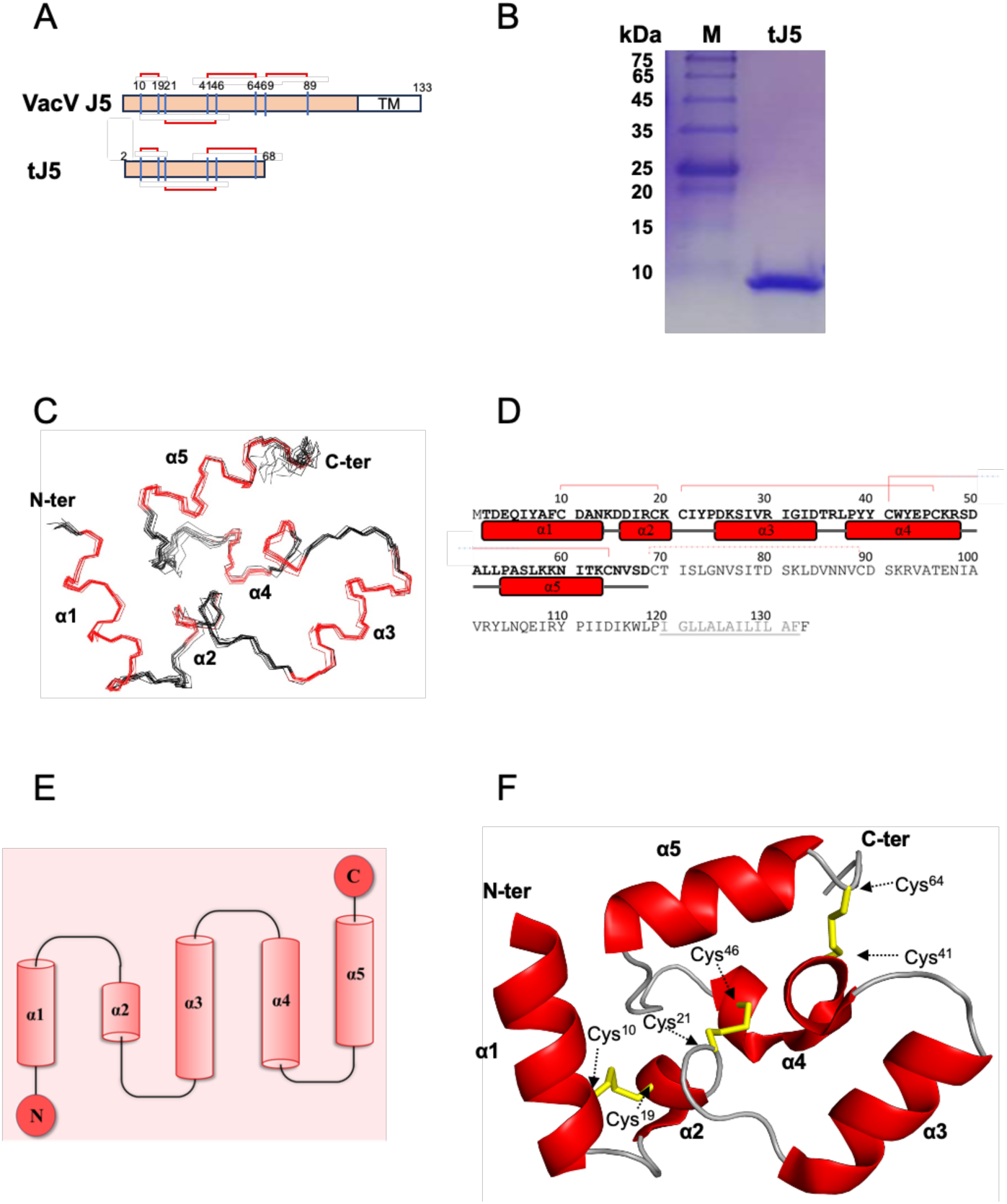
NMR structure determination of the vaccinia virus J5 ectodomain (tJ5, residues 2-68). (A) Schematic representation of the full-length J5 protein (residues 1-133) showing the transmembrane domain (TM; residues 111-133) and the truncated soluble construct (tJ5; residues 2-68) used for structural determination. (B) SDS-PAGE analysis of purified recombinant tJ5 protein stained with Coomassie blue. M, protein markers (kDa). (C) An ensemble of the 10 lowest-energy NMR structures of tJ5. (D) Secondary-structure organization of tJ5, consisting of five α-helices: α1 (residues 2-13), α2 (16–20), α3 (25–34), α4 (38–48), and α5 (54–63). Helices are represented as red cylinders; random coil regions are shown as gray lines. (E) Topological diagram of tJ5 generated using PDBsum. (F) The lowest-energy NMR structure of tJ5 at pH 6.5, as determined by NMR spectroscopy. Ribbon representation of the average NMR structure of tJ5, with α-helices are shown in red and three pairs of disulfide bonds, Cys10-Cys19, Cys21-Cys46 and Cys41-Cys64, highlighted in yellow.

To further elucidate the 3D structure of tJ5, we conducted multi-dimensional NMR experiments to gather information on NOE and dihedral angle restraints for protein structural determination. During structural refinement, we used an AMBER force field for energy minimization, incorporating molecules such as water and ions, in order to simulate the surrounding environment of tJ5 in solution. As a result, a high-resolution molecular structure with an average root mean square deviation (rmsd) of 0.5 Å was resolved, as summarized in Table 1. The NMR solution structure of tJ5 reveals five α-helices, namely α1 (T2-N13), α2 (D16-K20), α3 (D25-D34), α4 (P38-R48), and α5 (P54-K63), see Figure 1C-E. The overall structure of the ectodomain is stabilized by three disulfide bonds linking α1, α2, α4, and α5: C10-C19, C21-C46, and C41-C64 (Figure 1F). Notably, helices α1, α3, and α5 form a coplanar triangular ring structure, while both helices α2 and α4 extend off the plane in a concave conformation. These helices may interact with other fusion complex components in the EFC, playing an important role in virus infectivity.

### Functional mapping of VacV J5 protein in virus infectivity using recombinant viruses

To dissect the biochemical properties of J5 that contributes to vaccinia virus infection, a series of site-directed mutations were introduced based on surface exposure, charge distribution, and sequence conservation (Fig 2A). Exposed charged residues (D3, D11, K14) and conserved residues (P^38^YYWY^43^, S56, N65, N87, Y^103^Q^106^) were replaced with alanine or isoleucine. Multiple sequence alignment of J5 orthologs from representative orthopoxviruses and parapoxviruses (Fig. 2A) revealed high sequence identity within the N-terminal structural region (aa 2-68). Among them, AMV232 from *Amsacta moorei entomopoxvirus* showed the lowest similarity (27.9% identity, residues 2-68). Despite this divergence, AlphaFold-predicted models indicated that AMV232 adopts a protein fold similar to J5, with an RMSD of 0.662 Å across 57 Cα atoms. Minor deviations occurred at helix α3, while greater variation appeared in the unstructured loop preceding the transmembrane domain (Fig. 2B, right; Fig. S2A). The average pLDDT scores of full-length J5 and AMV232 were 80.7 and 76.5, respectively, with both proteins showing reduced confidence (<60) in the loop region before the transmembrane segment (Fig. S2B). To examine the contribution of this region, we swapped the J5 sequence between its two disulfide bonds with the corresponding AMV232 residues (Fig. 2B, left). The predicted structural similarity between J5 and AMV232 suggests these chimeric constructs likely maintain stable tertiary folds.

**Figure 2.**
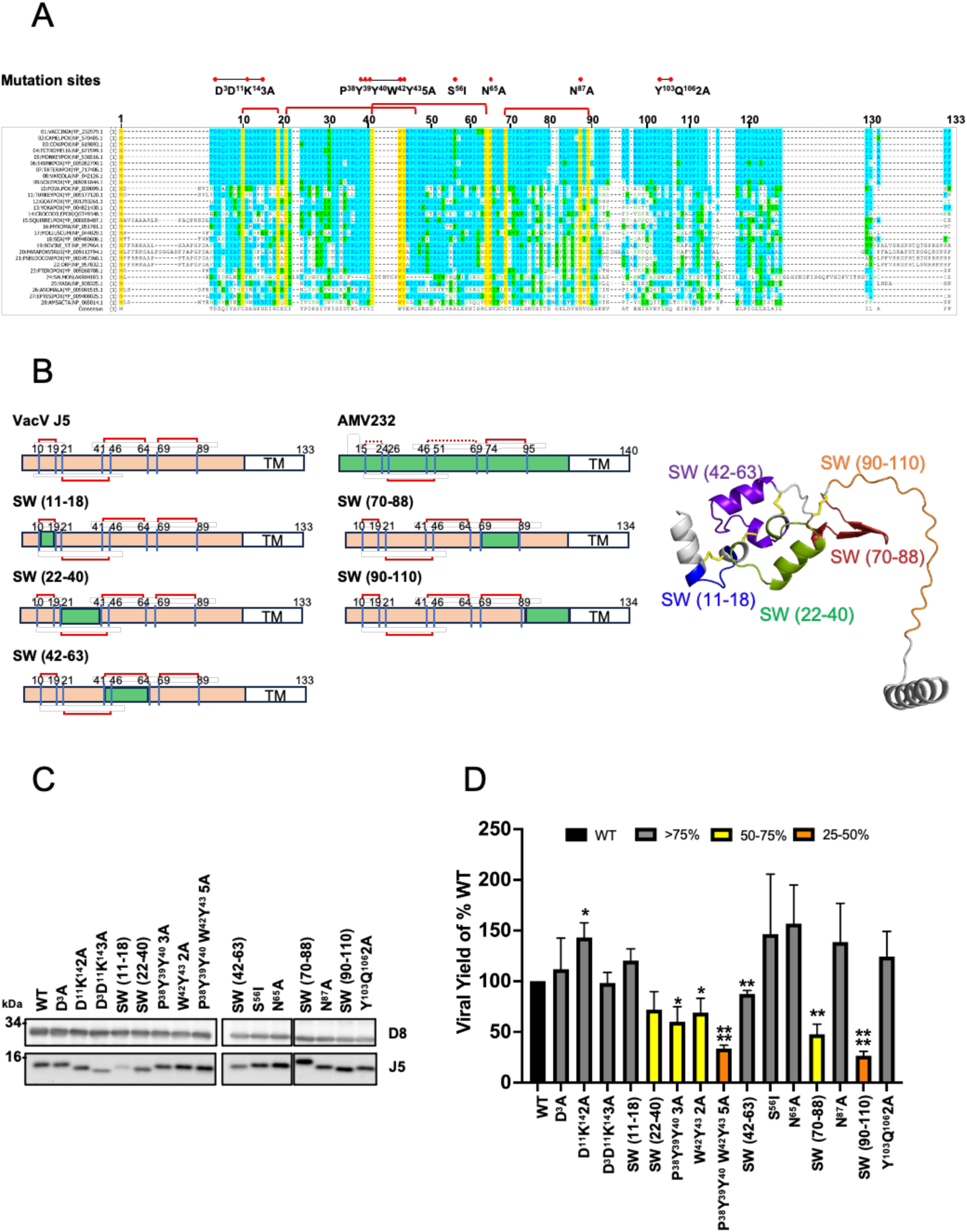
Functional mapping of the vaccinia virus J5 protein. (A) Multiple sequence alignment of J5 orthologues from representative poxviruses. Identical residues are shown in yellow (100% identity), moderately conserved residues in blue (>50%), and weakly conserved residues in green (>20%). Positions of the targeted mutations are indicated above the alignment. Predicted disulfide bonds are shown as red connecting lines. (B) Schematic representation of full-length vaccinia virus J5 (residues 1-133), AMV232 (residues 1-140), and the corresponding J5 swap mutants. The substituted regions (SW) are marked in colored boxes and mapped onto the predicted J5 three-dimensional model (right). (C) Immunoblot analysis of J5 and the structural protein D8 in infected cell lysates. (D) Virus yields of wild-type (WT) and mutant J5 recombinant viruses in BSC40 cells infected at a multiplicity of infection (MOI) of 5 PFU per cell and harvested 24 h post-infection for plaque assays. Infectivity of each mutant was normalized to that of WT J5. Data represent means ± standard deviations from three independent experiments. Statistical comparison of infectivity was performed between J5 WT and each mutant using Student’s *t*-test. \**P*<0.05; \*\**P*<0.01; \*\*\**P*<0.001; and \*\*\*\**P*<0.0001.

Because deletion of *J5L* is lethal for vaccinia virus replication (46), recombinant viruses expressing wild-type (WT) or mutant J5 were generated by homologous recombination. CV-1 cells were infected with WR-GS virus and transfected with individual J5 mutant plasmids containing the *gpt* selection cassette flanked by *J5L* homologous sequences. Recombinant viruses were isolated after three rounds of plaque purification in 1% agar under *gpt* selection, yielding a total of 16 recombinant viruses, including WT J5.

To assess the effect of each mutation on viral infectivity, BSC40 cells were infected at an MOI of 5 PFU/cell, and lysates harvested at 24 h post infection were analyzed by plaque assay and immunoblotting. Expression levels of J5 and the control protein D8 were comparable among most mutants and WT virus, except for the swap mutant 11-18, which showed a modest decrease in J5 expression (Fig. 2C). Infectivity of WT J5 was normalized to 100% (Fig. 2D). Most J5 mutants retained >75% infectivity (gray bars), while four mutants displayed moderate reductions (50-75%, yellow bars). Two mutants exhibited marked decreases (25-50%, orange bars). The most affected mutants mapped to residues 90-110, corresponding to the disordered loop preceding the transmembrane domain (Fig. 2B, right), and to the conserved P^38^YYCWY^43^ motif, a short helix shared among A16, G9, and J5 (P^38^YYCWY^43^ in J5; P^240^RECWD^245^ in G9; P^262^RVCWL^267^ in A16) (56). These regions, together with the swap mutant 22-40 (∼50% infectivity), appear critical for maintaining EFC function and viral entry efficiency.

### Defective J5 mutants with altered P^38^YYCWY^43^ helix and disordered C-terminal residues 90-110 display reduced fusogenic activity

BSC40 cells infected with equivalent amounts of each mutant virus particle were fixed and negatively stained at 24 h post-infection. TEM imaging (Fig. S3) showed typical mature virion morphology in both WT and mutant infections, consistent with previous findings that repression of EFC components does not affect virion morphogenesis or mature virion formation (11, 12, 16–18, 20–22, 33). We then investigated whether the infectivity loss of J5 mutants corresponds to their ability to trigger membrane fusion, using cell-cell fusion assays as previously described (32, 33). Lysates containing virus particles from the infectivity experiments were applied to GFP- and RFP-expressing HeLa cells mixed at 1:1 ratio for 1 h, followed by incubation in either neutral or acidic PBS to mimic endosomal acidification. Cells were then neutralized with growth medium and monitored for fusion events every 30 min for up to 3 h post-infection. Under neutral pH, little or no fusion was observed for WT or any mutant (Fig. 3A). In contrast, WT J5 induced robust syncytium formation following acid treatment (Fig. 3B). Fusion activity was quantified (Fig. 3C-D) as previously described (58). Acid-treated mutants carrying substitutions in the P^38^YYCWY^43^ helix or in the disordered C-terminal region (residues 90-110) exhibited significantly reduced fusion indices when compared to the WT virus (Fig. 3D), whereas most other mutants showed only mild reductions. Together, the data demonstrated that the conserved P^38^YYCWY^43^ motif and the flexible loop spanning residues 90-110 are critical determinants of J5-mediated membrane fusion.

**Figure 3.**
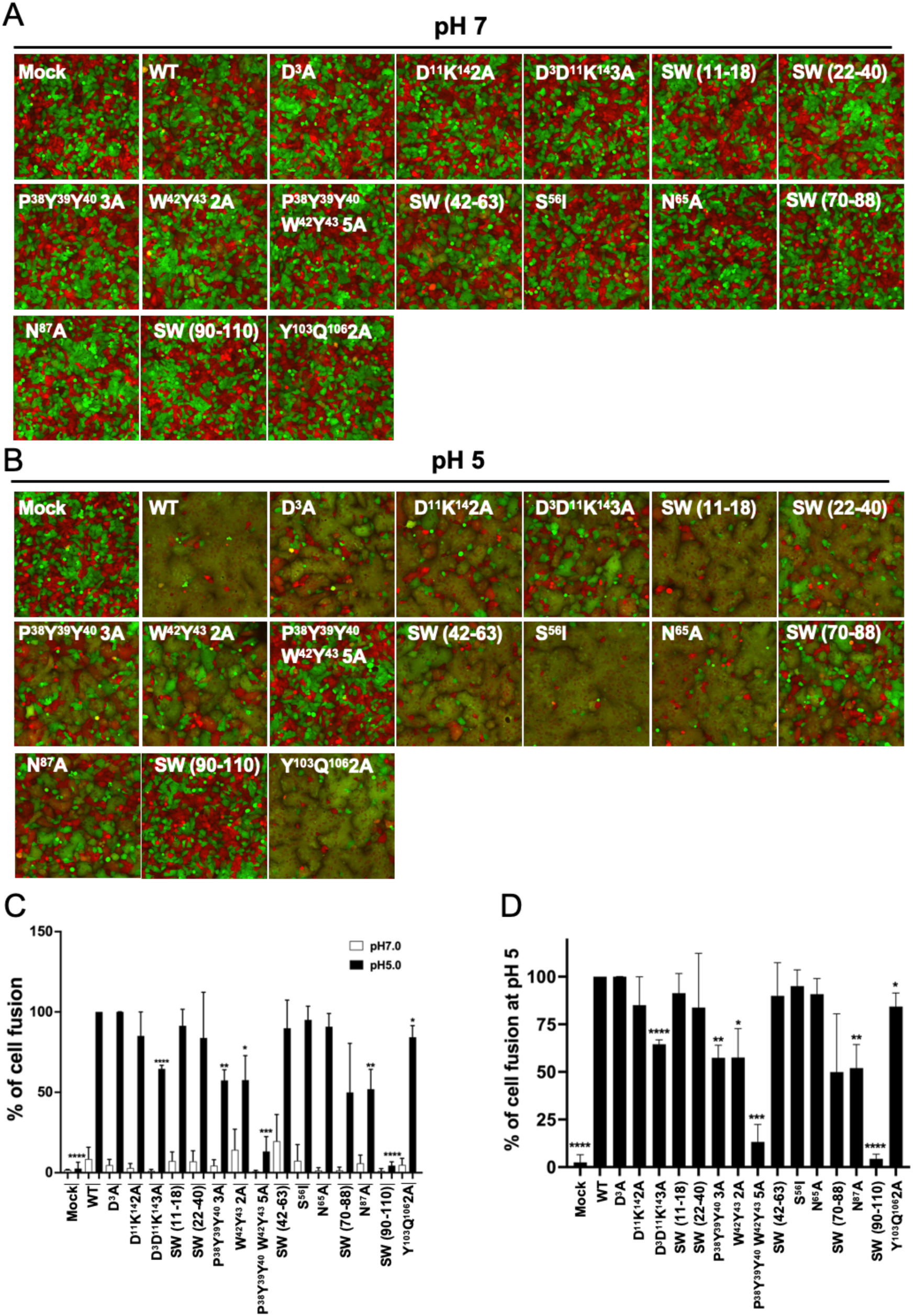
Cell-cell fusion assays of J5 mutants reveal impaired fusion activity at low pH. (A) HeLa cells expressing GFP or RFP were mixed at a 1:1 ratio and incubated with lysates containing wild-type (WT) or mutant J5 recombinant viruses under neutral pH conditions (pH 7). Fluorescence images were photographed at 3 h post-incubation. (B) Parallel fusion assays were performed as described in panel A, except that cells were incubated in acidic buffer (pH 5) for 3 min to mimic endosomal acidification as described in materials and methods. (C) Quantification of MV-triggered cell-cell fusion under neutral or acidic conditions. Images from three independent experiments were analyzed using Fiji software. Fusion was calculated as the percentage of the surface area of double-fluorescent cells relative to that of single-fluorescent cells. White and black bars represent fusion rates at pH 7 and pH 5, respectively. (D) Relative fusion activity of each mutant normalized to WT. Data represent means ± standard deviations from three independent experiments. Statistical comparison of fusion was performed between J5 WT and each mutant using two-tailed Student’s *t*-test. \**P*<0.05; \*\**P*<0.01; \*\*\**P*<0.001; and \*\*\*\**P*<0.0001.

### Residues 90-110 are required for incorporation of J5 into the EFC, while the P^38^YYCWY^43^ motif mediates interaction with peripheral components L1 and F9

To determine the basis of the fusion defects, we analyzed the assembly of the EFC in the two most impaired mutants, J5 ^90-110^ and J5 ^P38Y39Y40W42Y43-5A^, by coimmunoprecipitation. Anti-J5 antibody efficiently precipitated WT J5 together with all EFC components, whereas the J5 90-110 swap mutant failed to co-precipitate A16, G9, A28, H2, L1, and F9 (Fig. 4A, left). Reciprocal co-IPs using anti-G9 and anti-A28 antibodies showed that the G9-A16 and A28-H2 subcomplexes remained intact despite the absence of full complex assembly (Fig. 4A, middle and right). Thus, residues 90-110 are specifically required for incorporating J5 into the central trimer but are not needed for maintaining individual subcomplexes.

**Figure 4.**
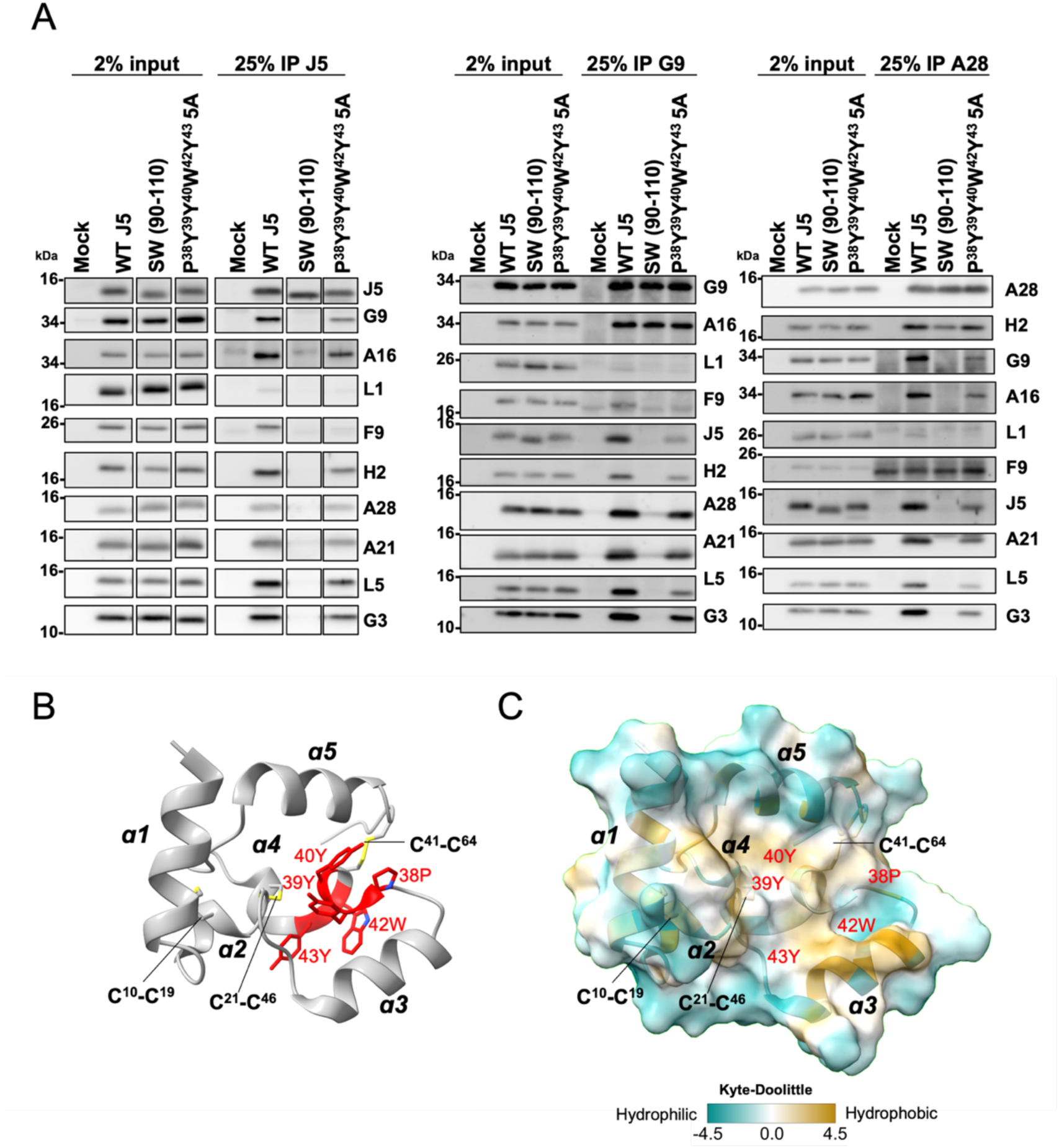
Defective J5 mutants disrupt assembly of the vaccinia virus EFC. (A) BSC40 cells were infected with wild-type (WT) or mutant J5 recombinant viruses. Soluble lysates collected 24 h post-infection were subjected to co-IP using antisera against J5 (left), G9 (middle), or A28 (right), followed by immunoblot analysis with antibodies against EFC components as indicated. The J5 SW(90–110) mutant failed to assemble a complete EFC but retained partial subcomplexes, whereas the J5^P38Y39Y40W42Y43-5A^ mutant exhibited impaired EFC formation, particularly affecting association with the peripheral protein F9. (B) Ribbon representation of tJ5 protein showing the P^38^YYCWY^43^ motif on helix α4 is centrally positioned among the surrounding α-helices. The side chains of P38, Y39, Y40, W42, and Y43 are shown in red. Secondary structure elements (α1-α5) and the three cysteine disulfide bonds (C10-C19, C21-C46, and C41-C46) are labeled. (C) Surface representation of the tJ5 protein colored according to the Kyte-Doolittle hydrophobicity scale, illustrating that the conserved the P^38^YYCWY^43^ motif harbors a hydrophobic core.

In contrast, the J5-5A mutant retained interactions with most EFC components but selectively lost binding to L1 and F9 (Fig. 4A, left). The tJ5 NMR structure places P^38^YYCWY^43^ within the α4 helix, whose packing is stabilized by three conserved disulfide bonds and aromatic stacking interactions (Fig. 4B, 4C) (56). This configuration forms a compact hydrophobic core on the N-terminal domain surface. Substituting these residues with alanine is predicted to destabilize this local fold, consistent with the selective disruption of L1/F9 association while preserving interactions with other EFC proteins. Together, these results reveal two functionally distinct elements within J5: the flexible 90-110 segment is essential for J5 incorporation into the EFC, whereas the P^38^YYCWY^43^ motif forms a localized structural module required for engaging the L1-F9 peripheral components.

### Structural mapping of J5 functional motifs and their roles in EFC assembly

To integrate the structural and functional data, we mapped the phenotypes of all J5 mutants onto the tJ5 NMR structure (Fig. 5A, top) and the AlphaFold2-predicted full-length J5 model (Fig. 5A, bottom). The most severe infectivity defects (<50% of WT) clustered in two regions: the N-terminal P^38^YYCWY^43^ motif and the flexible segment spanning residues 90-110. Mutants with intermediate phenotypes (∼50-75% of WT) localized adjacent to these same regions. In the NMR structure, the P^38^YYCWY^43^ motif forms an aromatic-cysteine cluster that stabilizes the α4 helix, whereas residues 90-110 map along the extended ectodomain-proximal helix in the full-length model. This analysis highlights two structurally distinct, mutation-sensitive elements within J5 (Fig. 5A)

**Figure 5.**
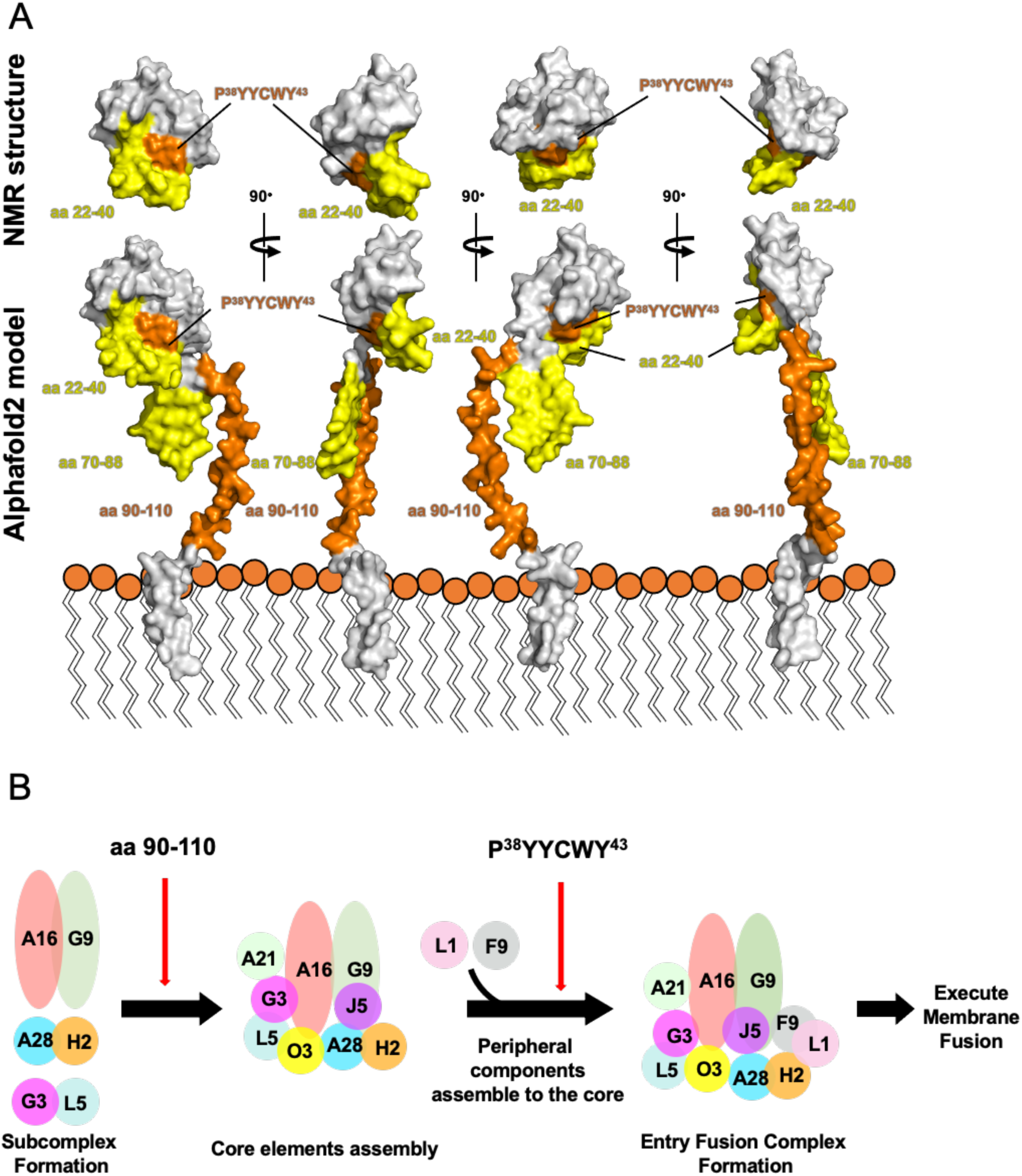
Summary of critical J5 residues and proposed model for EFC assembly. (A) Surface representation of the tJ5 NMR structure (top) and the AlphaFold2-predicted full-length J5 model (bottom), highlighting residues identified as critical for vaccinia virus infectivity. Regions in orange correspond to mutants showing <50% infectivity relative to wild type (WT), whereas regions in yellow indicate mutants retaining 50-75% infectivity. The locations of key functional regions, including residues 22-40, 70-88, and 90-110, are shown. (B) Proposed working model depicting the stepwise assembly of the vaccinia virus EFC. The disordered J5 loop (residues 90-110) mediates EFC assembly of EFC involving preformed subcomplexes, A16/G9, A28/H2, and G3/L5subcomplexes, while the conserved helix P^38^YYCWY^43^ facilitates recruitment of peripheral components F9 and L1. Proper integration of both regions enables the formation of the complete EFC required for membrane fusion.

Together, these findings define two mechanistically distinct roles for J5: residues 90-100 involves in a step to catalyze interaction of subcomplexes to form the 9-protein EFC core complements whereas the P^38^YYCWY^43^ motif provides a localized structural interface for EFC engaging L1/F9 peripheral components (Fig. 5B). These results establish a structure-function map for J5 and reveal how specific features of its ectodomain contribute to the ordered assembly and activation of the vaccinia virus entry fusion complex.

## Discussion

Virus entry, including receptor engagement and membrane fusion preceding genome release, represents a critical step in the infectious cycle and a prime target for antiviral intervention. Viral fusion proteins have historically been classified into three structural classes based on the organization of their fusion peptides or loops, which mediate membrane insertion (23, 59). However, the poxvirus entry fusion complex (EFC) remains unclassified, as its fusion machinery involves multiple viral proteins without sequence or structural similarity to canonical fusion motifs.

Bioinformatic analyses have shown that two conserved protein families, A16/G9/J5 and F9/L1, are found across diverse poxviruses and other large DNA viruses, implying an evolutionarily conserved fusion mechanism (45). Lipid-mixing assays further demonstrated that A28, L1, and L5 are dispensable for hemifusion (28), suggesting that fusogenic activity resides within the A16/G9/J5 orthologues. In this study, we determined the NMR structure of the J5 ectodomain (residues 2-68) and identified two regions critical for EFC function: the conserved P^38^YYCWY^43^ helix and the disordered C-terminal segment spanning residues 90-110.

Mutational analysis of 16 recombinant J5 viruses revealed that substitutions in other regions only modestly affected viral infectivity and cell-cell fusion, indicating that, despite low sequence conservation between vaccinia and AMV J5 proteins, most swapped J5 chimeras likely fold similarly to wild-type J5 and retain EFC assembly and fusion competence. In contrast, coimmunoprecipitation experiments demonstrated that the vaccinia J5 loop comprising residues 90-110 is indispensable for EFC formation through specific interactions with neighboring components, whereas replacement with the AMV equivalent loop abolished these contacts. The conserved P^38^YYCWY^43^ motif instead contributes primarily to the peripheral subunit F9 association.

Structural modeling of the chimeric J5 containing the AMV 90-110 loop using AlphaFold2, AlphaFold3, and RoseTTAFold yielded consistently low-confidence predictions, indicating intrinsic flexibility. Consistent with this, our recent cryo-EM reconstruction of the complete EFC (manuscript in revision) shows that J5 residues 90-110 form a flexible loop immediately preceding the transmembrane domain. This loop interlaces with neighboring loops from A16, G9, and A28, forming a membrane-proximal interaction network. Substitution of this region with the corresponding AMV232 sequence likely disrupts these loop-loop contacts, resulting in loss of EFC integrity and fusion activity.

Attempts to replace the vaccinia *J5L* open reading frame with full-length AMV232 under Eco-gpt selection were unsuccessful, consistent with prior reports that *J5L* is essential for virus replication (19, 46). Together, these findings underscore the crucial role of J5 in preserving the structural integrity of the EFC and in facilitating poxvirus membrane fusion.

All other ten EFC components have previously been characterized individually or within subcomplexes; the present study completes the structural map by defining the final component, J5. This work provides a structural and functional framework for understanding EFC organization and establishes the foundation for future mechanistic studies on poxvirus membrane fusion and its inhibition.

## Acknowledgments

We thank to Yae-Huei Liou and Wen-Li Pon at the IMB Imaging Core, Academia Sinica, and to Chen-Hsin Yu and Hsin-Nan Lin at the IMB Bioinformatics Core, Academia Sinica, and Shu-Yun Tung of the Genomics Core Facility at the IMB, Academia Sinica, for their assistance with imaging data analysis, bioinformatic analyses, and viral genome sequencing.

## Supplemental Figures

**Fig S1.**
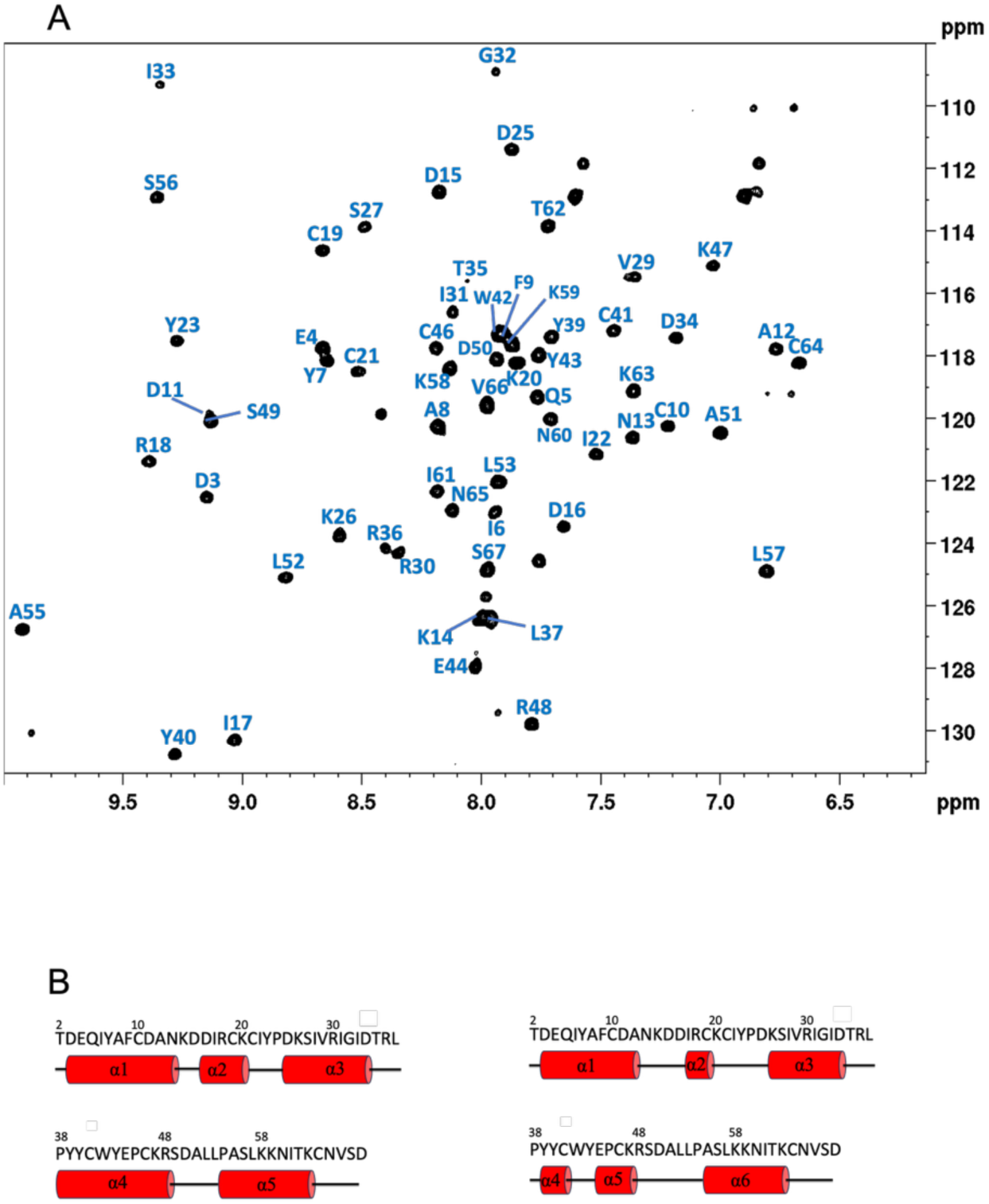
2D HSQC spectrum of vaccinia virus fusion component tJ5 and secondary structure predictions. (A) 2D ^1^H-^15^N HSQC spectrum of N-isotope-labeled tJ5 (0.2 mM) at pH 6.5. The spectrum was collected at 25 °C in an NMR buffer containing 10 mM phosphate and 137 mM NaCl and 2.7 mM KCl. The assigned residues are indicated using single-letter codes. (B) Secondary structure predictions of tJ5 by ^13^C_α_ chemical shift propensity (left panel) as well as AlphaFold2.0 (right).

**Fig S2.**
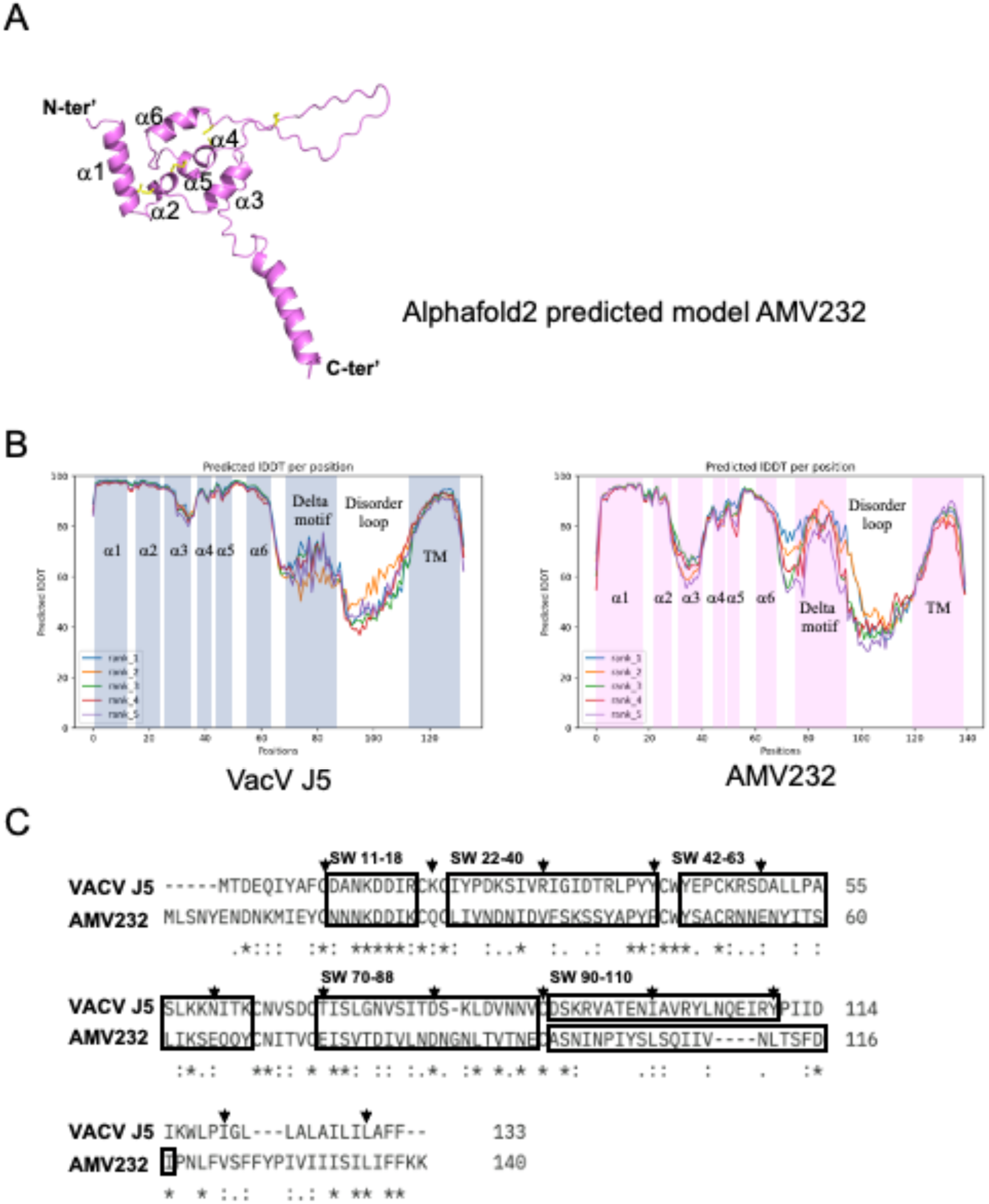
J5 and AMV23 shared conserved architecture and sequence, except the disorder region before the transmembrane region. (A) The AlphaFold 2 predicted AMV232 protein model. The ectodomain of AMV232 is similar to J5 (Figure 2B). (B) The pLDDT scores of AlphaFold 2 predicted full-length J5 and AMV232 protein. (C) The sequence alignment of vaccinia J5 and AMV232. The asterisk (*) represent the identical residues. The colon (:) imply the residues share similar biochemical properties. The dot (.) indicates the lower conservation between residues. The arrow marked the distance of 10 aa in the J5 sequence.

**Fig S3.**
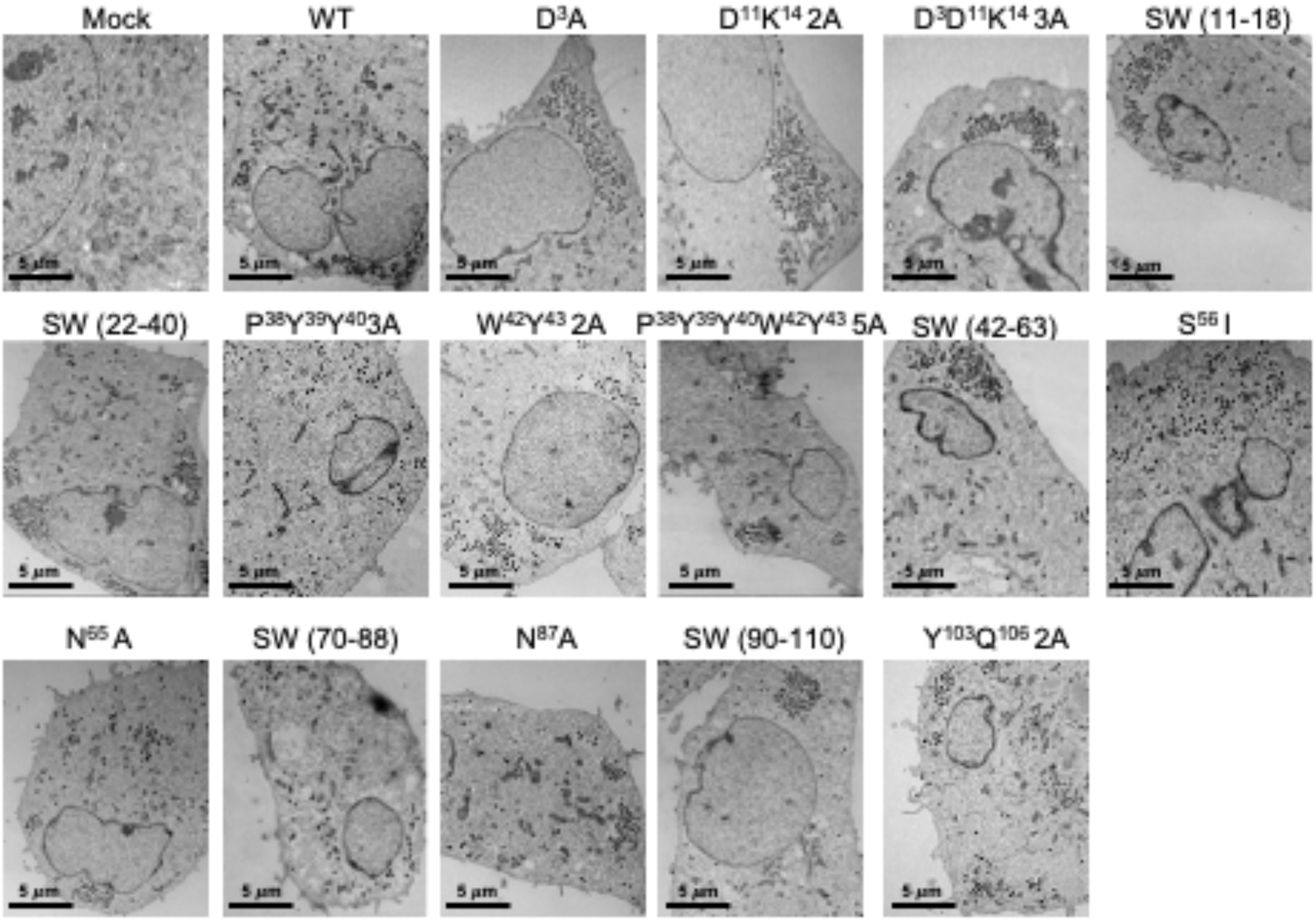
The normal morphogenesis of J5 WT and J5 mutant viruses. The BSC40 cells were infected with J5 WT and J5 mutant viruses for 24 h and processed for transmission electron microscopy as described in *Materials and Methods*. Scale bar, 5 μm.

**Fig S4.**
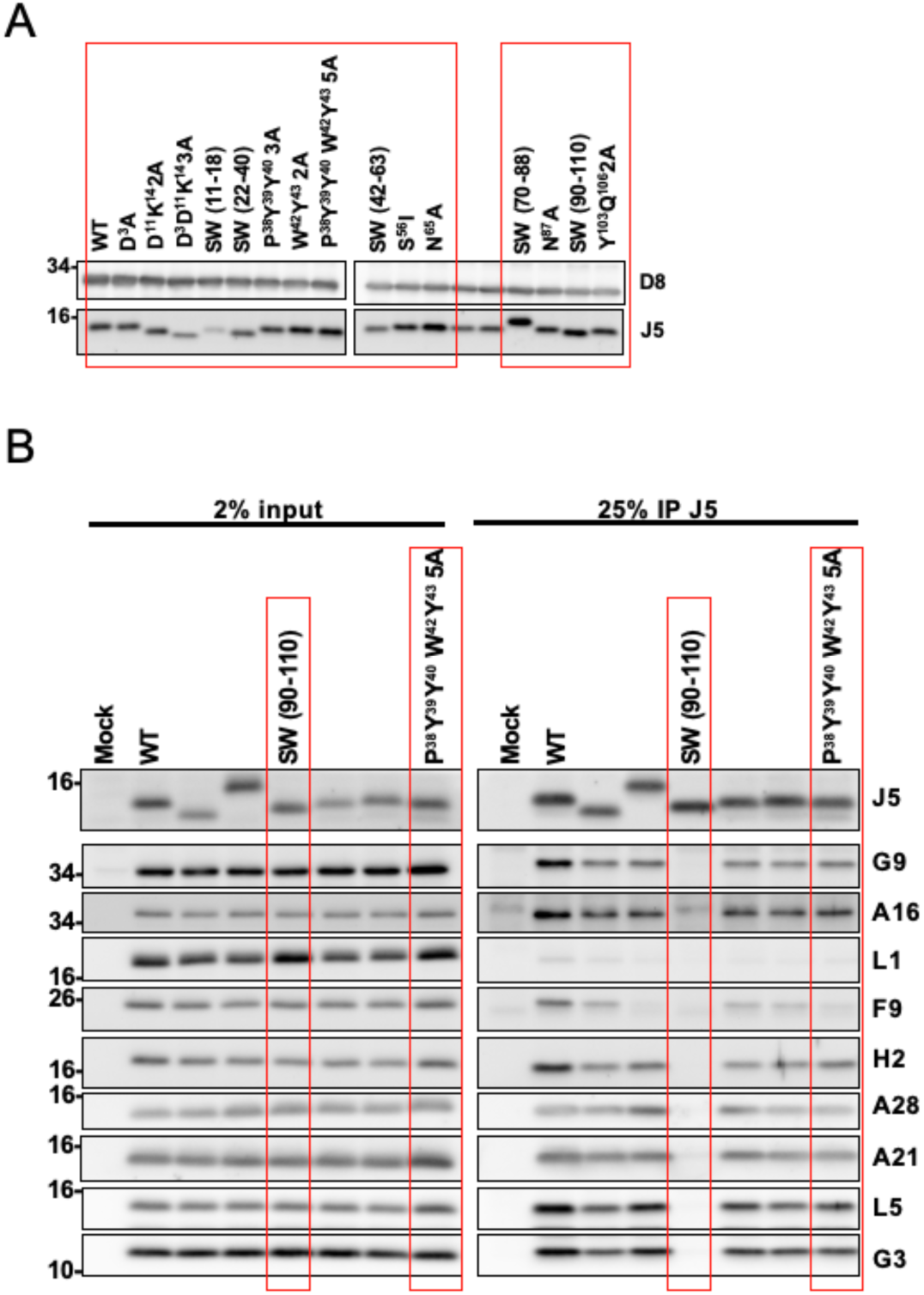
Uncropped immunoblot images referred to Figure 2C and 4A. The red boxes indicate the lanes shown in Figure 2C (A) and the left panel of Figure 4A (A).

